# DHODH inhibition enhances the efficacy of immune checkpoint blockade by increasing cancer cell antigen presentation

**DOI:** 10.1101/2023.04.03.535399

**Authors:** Nicholas J. Mullen, Surendra K. Shukla, Ravi Thakur, Sai Sundeep Kollala, Dezhen Wang, Nina Chaika, Juan F. Santana, William R. Miklavcic, Drew A. LaBreck, Jayapal Reddy Mallareddy, David H. Price, Amarnath Natarajan, Kamiya Mehla, David B. Sykes, Michael A. Hollingsworth, Pankaj K. Singh

## Abstract

Pyrimidine nucleotide biosynthesis is a druggable metabolic dependency of cancer cells, and chemotherapy agents targeting pyrimidine metabolism are the backbone of treatment for many cancers. Dihydroorotate dehydrogenase (DHODH) is an essential enzyme in the de novo pyrimidine biosynthesis pathway that can be targeted by clinically approved inhibitors. However, despite robust preclinical anticancer efficacy, DHODH inhibitors have shown limited single-agent activity in phase 1 and 2 clinical trials. Therefore, novel combination therapy strategies are necessary to realize the potential of these drugs. To search for therapeutic vulnerabilities induced by DHODH inhibition, we examined gene expression changes in cancer cells treated with the potent and selective DHODH inhibitor brequinar (BQ). This revealed that BQ treatment causes upregulation of antigen presentation pathway genes and cell surface MHC class I expression. Mechanistic studies showed that this effect is 1) strictly dependent on pyrimidine nucleotide depletion, 2) independent of canonical antigen presentation pathway transcriptional regulators, and 3) mediated by RNA polymerase II elongation control by positive transcription elongation factor B (P-TEFb). Furthermore, BQ showed impressive single-agent efficacy in the immunocompetent B16F10 melanoma model, and combination treatment with BQ and dual immune checkpoint blockade (anti-CTLA-4 plus anti-PD-1) significantly prolonged mouse survival compared to either therapy alone. Our results have important implications for the clinical development of DHODH inhibitors and provide a rationale for combination therapy with BQ and immune checkpoint blockade.

## Introduction

Deranged cellular metabolism is a universal feature of cancer cells ^1,2^. One particularly cancer-essential metabolic aberration is the hyperactive synthesis and utilization of nucleotide triphosphates; this phenotype is a critical driver of cancer cell malignant behaviors, including uncontrolled proliferation, evasion of the host immune response, metastasis to distant organs, and resistance to antineoplastic therapy ^3^. The de novo pyrimidine biosynthesis pathway, which generates pyrimidine nucleotides from aspartate and glutamine, is consistently hyperactive in cancer cells and druggable by clinically approved inhibitors ^4^. Dihydroorotate dehydrogenase (DHODH) catalyzes the fourth step in this pathway and is essential for de novo pyrimidine synthesis. DHODH inhibitors have shown robust preclinical anticancer activity across diverse cancer types ^5-14^ and have recently entered clinical trials for multiple hematologic cancers (NCT04609826 and NCT02509052). Although there is a vast literature on DHODH inhibitors dating back to the early 1990s, and despite the “rediscovery” of DHODH in recent years as a critical cancer cell metabolic dependency, important questions about the cellular response to DHODH inhibition remain unanswered.

While combination chemotherapy is highly effective and potentially curative against certain cancers (e.g. Hodgkin lymphoma, testicular cancer, childhood leukemia, and others), many common malignancies are refractory to chemotherapy (e.g. lung cancer, pancreatic cancer, colorectal cancer, etc.) ^15^. In some chemotherapy-refractory cancers (most prominently melanoma, mismatch repair deficient colorectal cancer, bladder cancer, and non-small cell lung cancer), immunotherapeutic strategies have demonstrated strong efficacy and led to durable remissions in a subset of patients ^16^. The efficacy of immunotherapy agents is dependent on multiple factors, including tumor antigen presentation, limited immune cells in the tumor milieu, and T-cell activation status ^17,18^. Adoptive cell therapies and immune checkpoint blockade (ICB) can address the issues of limited immune cell recruitment into tumors and limited T-cell activation, respectively. However, adequate antigen presentation by tumor cells is still required for immunotherapy efficacy, which relies on T-cell-mediated adaptive immunity.

The antigen presentation pathway (APP) mediates the presentation of endogenous peptide antigens to CD8 T-cells via MHC class I (MHC-I). This pathway entails the degradation of cellular proteins into small peptides by the proteasome, the import of these peptides into the endoplasmic reticulum by transporter associated with antigen presentation proteins (*TAP1* and *TAP2*), and the loading of these peptides into the MHC-I complex, which consists of a heavy chain (encoded by *HLA-A, HLA-B*, or *HLA-C*) and a light chain (encoded by *B2M*) ^19^. APP genes are often downregulated in cancer cells, and this impedes the recognition of immunogenic MHC-I restricted cancer cell antigens by infiltrating T-cells ^20^. Antigen presentation and T-cell recognition are crucial for T-cell-mediated killing of cancer cells ^21-23^, and forced MHC-I expression enhances immunotherapy efficacy in preclinical models ^24-27^. Furthermore, high tumoral expression of MHC-I, MHC-II, and other APP genes correlates with better overall survival in patients with melanoma treated with ICB therapies ^28-31.^

While previous reports have shown that pyrimidine nucleotide depletion triggers the expression of innate immunity-related genes and induces an interferon-like response ^32-34^, the role of pyrimidine starvation in antigen presentation has not been reported. Herein, we report that DHODH inhibition induces the robust upregulation of APP genes and increases tumor cell antigen presentation via MHC-I. We further explored the mechanism and functional consequences of DHODH inhibitor-mediated APP induction in cancer.

## Results

### Brequinar induces upregulation of MHC-I and antigen presentation pathway genes

We examined gene expression changes following transient or prolonged DHODH inhibition by culturing human pancreatic ductal adenocarcinoma cell lines S2-013 and CFPAC-1 in the presence of BQ at two different doses for 16 hours and for a two-week duration (Fig 1A). Gene set enrichment analysis (GSEA) using Hallmark and KEGG gene sets from MSigDB ^35,36^ revealed 17 gene sets that were significantly upregulated (FDR q < 0.25) across both cell lines following two-week BQ exposure (Fig 1B). Twelve of these gene sets (highlighted in purple) are ontologically related to antigen presentation and contain MHC class I, MHC class II, and/or APP genes such as *TAP1* in the leading edge. Certain gene sets, such as allograft rejection (KEGG), graft versus host disease (KEGG), and antigen processing and presentation (KEGG) are composed almost entirely of APP genes (Fig 1C). Heatmap analysis showed that APP genes were robustly upregulated in a dose- and duration-dependent manner in CFPAC-1 (Fig 1D) and S2-013 (Fig S1A) cells. The effect size was generally smaller for S2-013 cells, likely because they are resistant to DHODH inhibition due to efficient nucleoside salvage, as we previously reported ^37^. Publicly available RNA-seq data from human A375 melanoma cells treated with the clinically approved DHODH inhibitor teriflunomide ^38^ corroborated our findings, as teriflunomide caused a rapid (within 12 hours) and duration-dependent increase in MHC-I/II and APP transcript levels (Fig 1E).

**Figure 1:**
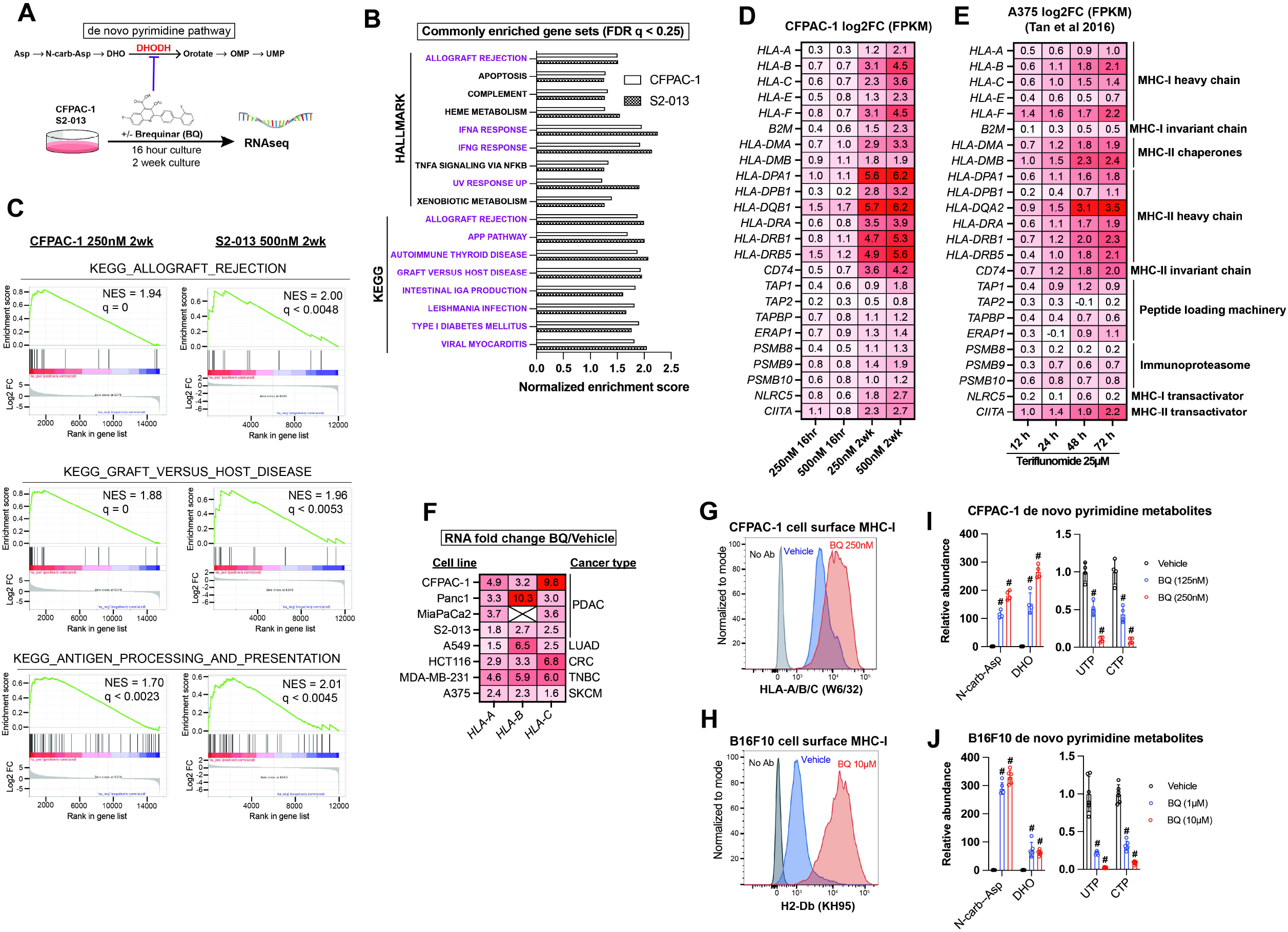
Brequinar induces mRNA expression of antigen presentation pathway genes and upregulates cell surface MHC-I in diverse cancer cell lines. **A)** Schematic of RNA sequencing experiment for panels C-D, with de novo pyrimidine pathway shown to highlight the role of DHODH **B)** Normalized enrichment scores for gene sets commonly enriched (FDR q < 0.25) in S2-013 (500nM) and CFPAC-1 (250nM) cells following two-week BQ treatment, as assessed by gene set enrichment analysis. **C)** GSEA plots for indicated gene sets following two-week BQ treatment of CFPAC-1 (left) or S2-013 (right) cells at the indicated doses **D)** Heatmap showing log2 fold change mRNA expression of APP genes in CFPAC-1 cells treated with BQ for indicated dose and duration. **E)** Heatmap showing log2 fold change mRNA expression for APP genes in A375 melanoma cells treated with the DHODH inhibitor teriflunomide (25μM) for indicated durations, data extracted from ^38^. **F)** RT-qPCR quantification of *HLA-A, HLA-B*, and *HLA-C* mRNA levels in cancer cell lines after 24-hour BQ treatment. Numbers represent fold change relative to vehicle control for each cell line. Data are representative of at least 3 independent experiments. *HLA-B* was not detectable in MiaPaCa2 cells. **G-H)** Flow cytometry analysis of cell surface MHC-I in live CFPAC-1 (G) or B16F10 (H) cells following 10-day treatment with BQ (250nM for CFPAC-1 and 10μM for B16F10). **I-J)** LC-MS/MS metabolomics quantification of de novo pyrimidine pathway metabolites in CFPAC-1 (I) or B16F10 (J) cells following 8-hour BQ treatment at indicated doses. Data represent mean +/-SEM of four (CFPAC-1) or six (B16F10) biological replicates. # indicates p < 0.0001 by two-way ANOVA with Bonferonni’s post-comparison test.

We validated these gene expression changes in CFPAC-1 cells by RT-qPCR (Fig S1B) and then performed RT-qPCR to assess the mRNA levels of genes coding for MHC-I across a panel of human cancer cell lines treated with BQ for 24 hours (Fig 1F). This confirmed that MHC-I heavy chain transcripts (*HLA-A, HLA-B*, and *HLA-C*) are consistently upregulated in response to BQ across diverse cancer types (Fig 1F). To optimize conditions for *in vivo* studies, we tested the long-term response and observed that two-week BQ treatment of B16F10 murine melanoma cells also caused dramatic APP gene upregulation (Fig S1C). Flow cytometry confirmed a marked increase in cell surface MHC-I levels in nonpermeabilized live CFPAC-1 (Fig 1G) and B16F10 (Fig 1H) cells following a two-week BQ treatment, confirming that transcriptional upregulation of APP genes results in greater cell surface antigen presentation.

In parallel, we confirmed pyrimidine nucleotide depletion upon treatment with BQ at different doses by performing metabolomics analysis of CFPAC-1 and B16F10 cells following BQ treatment. The results demonstrated a rapid (8-hour treatment) and dose-dependent accumulation of dihydroorotate and N-carbamoyl-aspartate (upstream of DHODH) as well as depletion of pyrimidine nucleotides UTP and CTP (Fig 1I-J) and other pyrimidine species (Fig S1D-E). These results confirm that on-target DHODH inhibition and resultant pyrimidine nucleotide depletion correlates with the transcriptional induction of APP genes and enhanced antigen presentation via MHC-I.

### BQ-mediated APP induction depends on pyrimidine nucleotide depletion

To confirm that BQ-or teriflunomide-mediated APP induction was specifically caused by DHODH inhibition (i.e., on-target effect), we asked whether the effect could be reversed by restoring pyrimidine nucleotides. As we previously observed ^37^, media supplementation with uridine rescued cell viability (Fig 2A) and pyrimidine levels (Fig 2B) following BQ treatment and partially rescued viability following teriflunomide treatment (Fig S2A). Uridine supplementation likewise blocked mRNA induction of MHC-I transcripts (*H2-Db, H2-Kb*, and *B2m*), as well as *Nlrc5* (a major MHC-I transcriptional coactivator) and *Tap1* (required for peptide import into the ER, a key step in MHC-I antigen presentation) by BQ (Fig 2C) or teriflunomide (Fig S2B), while uridine alone had no effect (Fig S2C). This same phenotype was observed in HCT116 human colorectal cancer cells (Fig S2D). Concordantly, cell surface MHC-I upregulation by BQ or teriflunomide (24-hour treatment) was abrogated by uridine supplementation (Fig 2D), while uridine alone again had no effect (Fig S2E). These results demonstrate that DHODH inhibitor-mediated APP induction is caused by pyrimidine nucleotide depletion.

**Figure 2:**
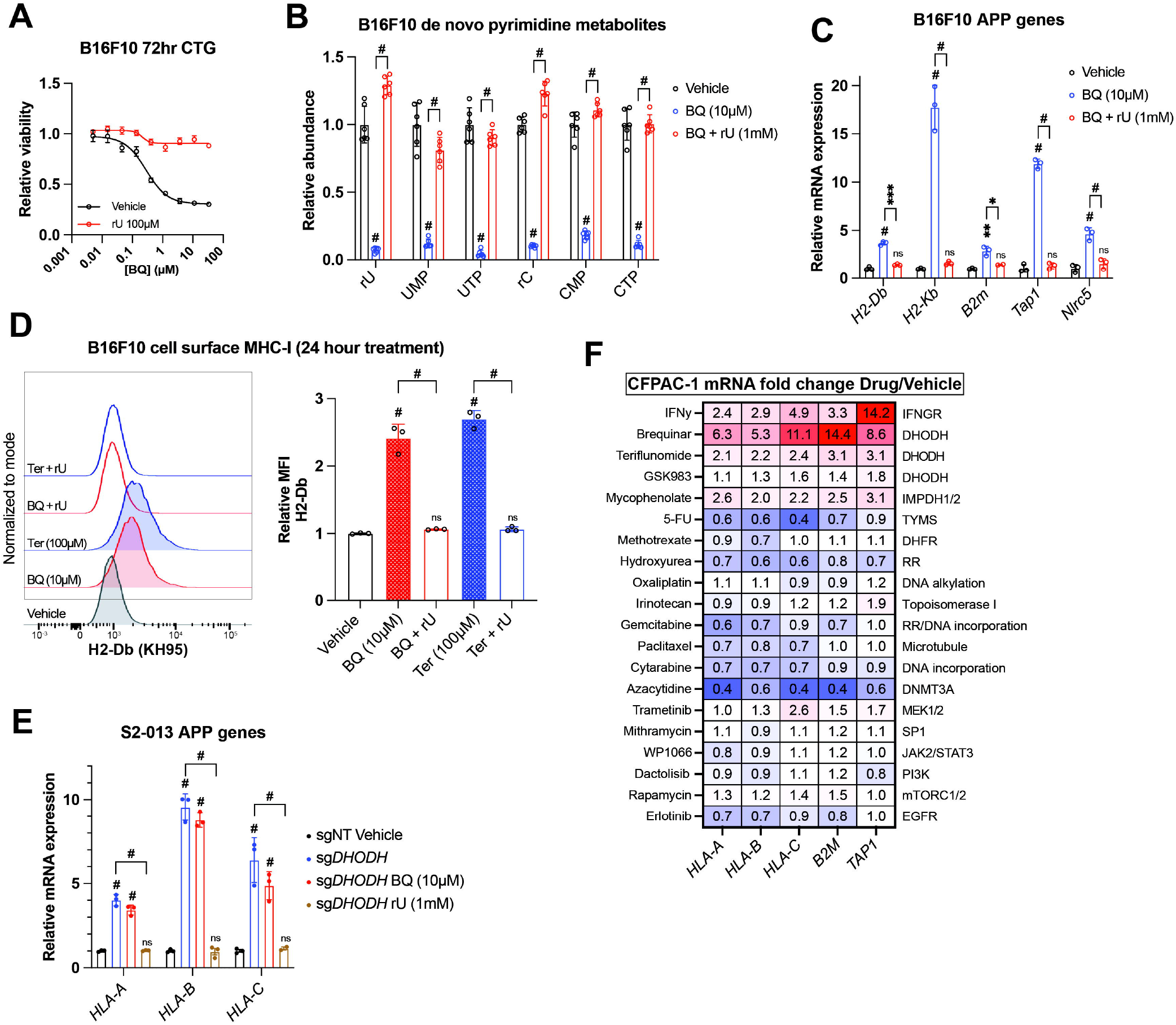
BQ-mediated APP induction requires pyrimidine nucleotide depletion. **A)** Dose-response cell viability experiment in B16F10 cells treated with BQ +/-uridine (100μM) for 72 hours. Data represent mean +/-SEM of three biological replicates. One representative result of three independent experiments is shown. **B)** Quantification of pyrimidine metabolites following 24-hour treatment of B16F10 cells with vehicle, BQ (10μM), or BQ + uridine (1mM). Data represent mean +/-SEM of six biological replicates. # indicates p < 0.0001 by two-way ANOVA with Bonferonni post-comparison test. **C)** RT-qPCR of indicated genes in B16F10 cells following 24-hour treatment with BQ (10μM) +/-uridine (1mM). Data represent mean +/-SD of three technical replicates. One representative result of three independent experiments is shown. **D)** Left: flow cytometry analysis of cell surface MHC-I (H2-Db) on live B16F10 cells following 24-hour treatment with indicated agents (BQ 10μM, teriflunomide 100μM, uridine 1mM). Right: quantification of H2-Db mean fluorescence intensity normalized to vehicle control. Data represent mean +/-SEM of three independent experiments. # indicates p < 0.0001 with two-way ANOVA with Bonferroni post-comparison test. **E)** RT-qPCR analysis of indicated genes in S2-013 cells with DHODH knockout (sgDHODH) or non-targeting control vector (sgNT) treated with indicated agents for 72 hours. Data represent mean +/-SD of four determinations. One representative result of three independent experiments is shown. **F)** RT-qPCR analysis of indicated genes in CFPAC-1 cells following 72-hour treatment with indicated agents. Numbers in the heatmap represent mean fold change versus vehicle control with four determinations.

To further validate this finding, we assessed MHC-I heavy chain mRNA levels in S2-013 cells with DHODH deletion (sgDHODH). We have previously demonstrated that these cells require exogenous uridine for viability and experience profound pyrimidine depletion (>95% depletion of UTP and CTP) after 8-hour incubation in nucleoside-free media ^37^. After growing these cells with supplemented uridine (1 mM), we withdrew exogenous nucleosides by changing to new media containing 10% dialyzed FBS. After 72-hour exposure to nucleoside-free media, sgDHODH cells upregulated *HLA-A, HLA-B*, and *HLA-C*, and this was reversed by adding back uridine (Fig 2E). Importantly, treatment with BQ did not further increase MHC-I mRNA expression (Fig 2E, compare blue and red bars). Together with our other data, these results indicate that BQ-mediated APP induction is an on-target phenomenon with respect to DHODH inhibition.

Since uridine addback rescued BQ- and teriflunomide-mediated loss of viability (Fig 2A, S2A), we queried whether BQ-mediated APP induction was caused by pyrimidine depletion *per se*, or if it was the result of some nonspecific downstream consequence of pyrimidine starvation, such as DNA damage or loss of cell viability. To address this, we screened a panel of genotoxic chemotherapy agents and small molecule inhibitors for their ability to induce APP genes following 72-hour exposure at previously determined cytotoxic doses in CFPAC-1 cells (Fig 2F). Besides interferon gamma (a positive control), BQ, teriflunomide, and GSK983 (another DHODH inhibitor), the only agent that induced APP gene transcription in this assay was mycophenolate, a clinically approved inhibitor of the *de novo* GTP synthesis enzymes inosine monophosphate dehydrogenase 1 and 2 (IMPDH1/2). The effect of mycophenolate on APP gene expression was subsequently validated in B16F10 cells (Fig S2F), demonstrating that either purine or pyrimidine nucleotide depletion can induce cancer cell APP mRNA expression *in vitro*.

The other drugs screened included nucleotide synthesis inhibitors (5-fluorouracil, methotrexate, gemcitabine, and hydroxyurea), DNA damage inducers (oxaliplatin, irinotecan, and cytarabine), a microtubule targeting drug (paclitaxel), a DNA methylation inhibitor (azacytidine), and other small molecule inhibitors (Fig 2F). While we cannot rule out the possibility that these agents induce APP transcription in other cell lines or under other dose/duration conditions, the inertness of these compounds (with respect to APP gene expression) in our screen suggests that BQ-mediated APP induction in CFPAC-1 cells is not a general phenomenon that occurs downstream of DNA damage or some other response to therapy-induced stress.

### BQ-mediated APP induction does not depend on canonical APP transcriptional regulators

To elucidate the molecular pathway leading to APP induction downstream of pyrimidine depletion, we extended our findings to HEK-293T cells, which also display rapid (within 4 hours) transcriptional induction of MHC-I upon BQ treatment (Fig S3A). Reasoning that the mechanism of this phenomenon in HEK-293T cells is less likely to involve idiosyncratic genetic aberrations than in cancer cell lines, we chose to conduct our initial mechanistic studies in this system and then extend our findings to cancer cell lines if possible.

We used a candidate-based chemical biology screening approach to ask if drugs targeting suspected pathways might block BQ-mediated APP induction in HEK-293T cells. We first interrogated pathways that are known to control MHC/APP expression, including IFN-JAK-STAT ^39^, NF-ĸB ^27,40^, and cGAS-STING-TBK1 ^41.^ Neither ruxolitinib (a JAK1/2 inhibitor with activity against STAT3) nor GSK8612 (a TBK1 inhibitor) ^42^, nor TPCA-1 (an IKK2 inhibitor) ^43^ abrogated BQ-mediated APP induction (Fig 3A), despite blocking APP induction downstream of poly(dA:dT) and interferon gamma (Fig S3B) as expected. This indicates that these canonical regulators of MHC/APP expression are dispensable for APP induction downstream of DHODH inhibition.

**Figure 3:**
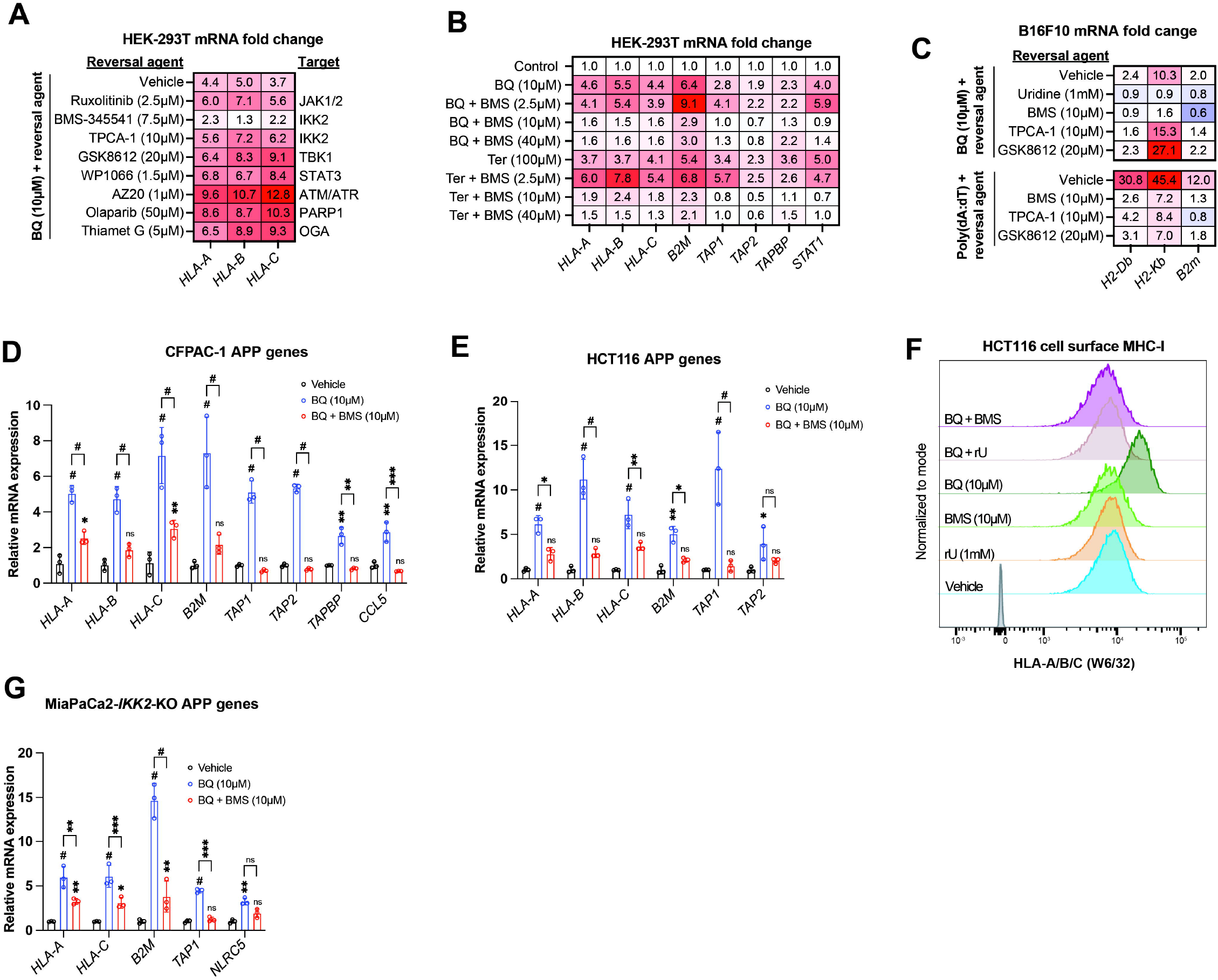
IKK2 inhibitor BMS-345541 abrogates BQ-mediated APP induction in an IKK2-independent manner. **A-B)** HEK-293T cells were treated with indicated agents for 24 hours and then subjected to RT-qPCR analysis for indicated genes. Numbers in the heatmap represent mean of four determinations. **C-E, G)** B16F10 (C), CFPAC-1 (D), HCT116 (E) or MiaPaCa2-IKK2-KO (G) cells were treated with indicated agents for 24 hours and subjected to RT-qPCR analysis of indicated genes. Data in D,E, and G represent mean +/-SD of three independent experiments. * indicates p < 0.05, ** p < 0.01, *** p < 0.001, and # p < 0.0001 with two-way ANOVA with Bonferroni post-comparison test. For C, numbers in the heatmap represent mean fold change versus vehicle with three determinations. Representative result of three independent experiments is shown. **F)** Flow cytometry analysis of cell surface MHC-I in HCT116 cells treated with indicated agents for 24 hours.

Interestingly, the IKK2 inhibitor BMS-345541 ^44^ mostly abrogated BQ-mediated APP induction (Fig 3A). BMS-345541 effectively blocked BQ- and Ter-mediated APP induction at concentrations of 10 μM and 40 μM, but not 2.5 μM (Fig 3B). The effect of BMS-345541 was confirmed in B16F10 (Fig 3C), CFPAC-1 (Fig 3D), and HCT116 (Fig 3E) cells. Furthermore, BQ treatment (24 hours) of HCT116 cells caused increased cell surface expression of MHC-I, which could be reversed by either uridine supplementation or by treatment with BMS-345541; neither uridine nor BMS-345541 alone affected cell surface MHC-I expression (Fig 3F).

Given that TPCA-1 (an established IKK2 inhibitor ^43^) did not block BQ-mediated APP induction (Fig 3A, 3C), we suspected that this effect of BMS-345541 was independent of IKK2. To test this, we used previously reported MiaPaCa2 cells with CRISPR-Cas9 deletion of *IKK2* (MiaPaCa2-*IKK2*-KO) ^45^. Increased APP mRNA expression was observed upon BQ, teriflunomide, or GSK983 treatment (all DHODH inhibitors) of either wild-type or *IKK2*-KO MiaPaCa2 cells (Fig S3C). However, while TNF-alpha stimulation induced APP and *CCL5* (a canonical NF-ĸB target gene downstream of TNF-alpha ^46^) expression in wild-type cells, this was not observed in *IKK2*-KO cells, as expected (Fig S3C, far right). Finally, BQ-mediated APP induction in *IKK2*-KO cells was significantly reversed with concurrent BMS-345541 treatment (Fig 3G). Together, these results demonstrate that IKK2 is dispensable for BQ-mediated APP induction and that the observed reversal effect of BMS-345541 is independent of IKK2.

### Nucleotide starvation induces APP transcription in a P-TEFb-dependent manner

To further investigate the mechanism by which BMS-345541 blocks APP induction downstream of pyrimidine starvation, we leveraged publicly available data on the target profile of BMS-345541 and other agents tested in the cell-free KINOMEscan assay ^47^. BMS-345541 reproducibly bound more than 20 kinases, with dissociation constants (k_d_) ranging from 130-8100 nM (Fig 4A). We prioritized potential targets with a k_d_ in the low micromolar range, given that 2.5 μM BMS-345541 did not block BQ-mediated APP induction in our previous experiments, and the effect seemed to be maximal at 10 μM, with no significant increase in the magnitude of the effect between 10 μM and 40 μM (Fig 3B). Additionally, we prioritized targets that were >50% inhibited with 10 μM BMS-345541 treatment. These two conditions correspond to the upper left quadrant of Fig 4A.

**Figure 4:**
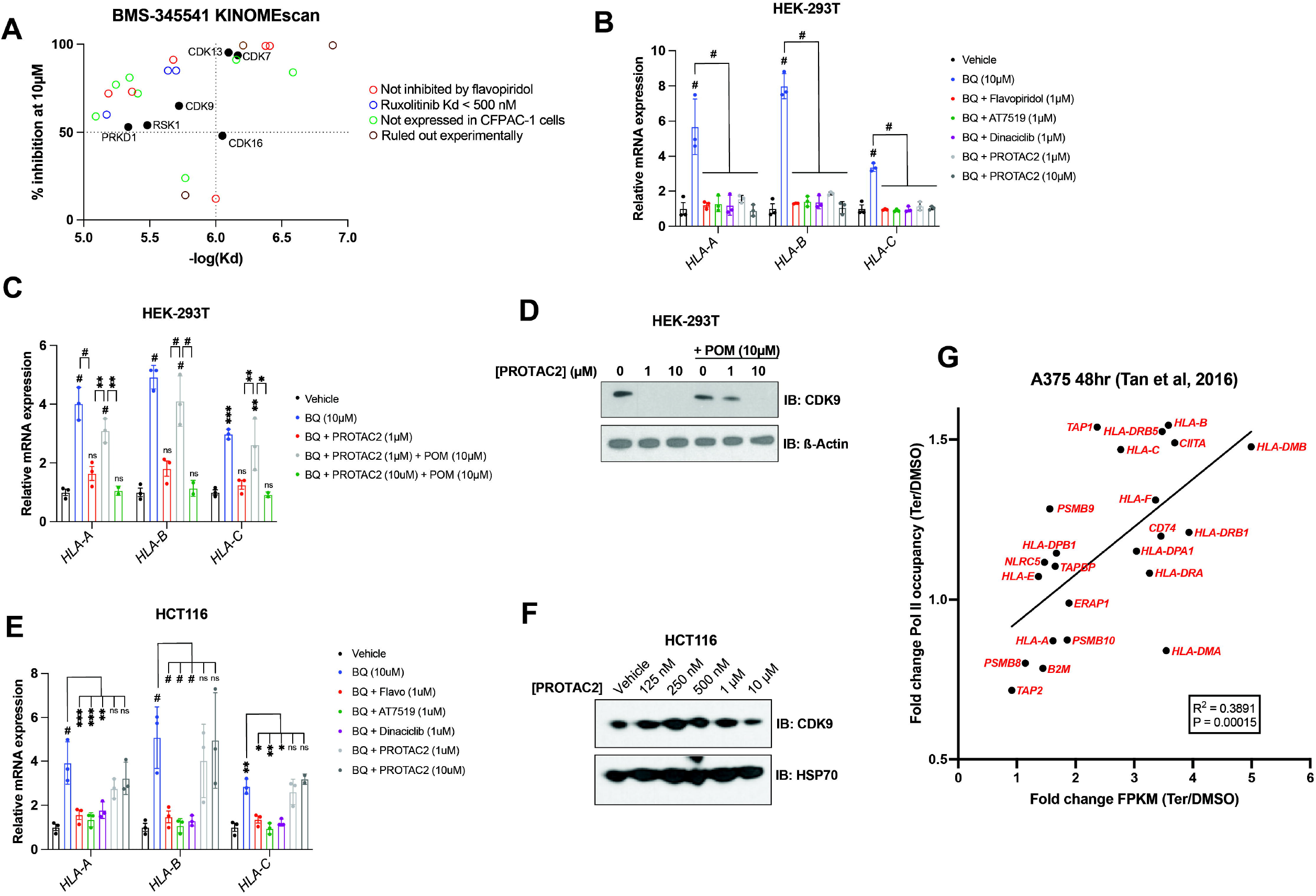
P-TEFb inhibitor flavopiridol abrogates APP induction downstream of nucleotide depletion. **A)** Plot of percent inhibition (10μM treatment) vs -log(dissociation constant) for kinases bound by BMS-345541 in KINOMEscan assays ^47^. Each data point represents an individual kinase. **B-C)** RT-qPCR analysis for indicated genes in HEK-293T cells treated with indicated agents for 24 hours. Data represent mean +/-SD of three independent experiments. * indicates p < 0.05, ** indicates p < 0.01, *** indicates p < 0.001, and # indicates p < 0.0001 by two-way ANOVA with Bonferonni’s post-comparison test **D)** Western blot analysis for CDK9 performed on HEK-293T cells treated with CDK9-targeted PROTAC (PROTAC2) and/or pomalidomide (POM) for 24 hours. Beta actin was used as a loading control. **E)** RT-qPCR analysis of indicated genes after 24-hour treatment with indicated agents. Data represent mean +/-SD of three independent experiments. **F)** Western blot analysis for CDK9 performed on HCT116 cells treated with the indicated concentrations of PROTAC2 for 24 hours. Heat shock protein 70 (HSP70) was used as a loading control **G)** Linear regression analysis of fold change (teriflunomide/DMSO) in Pol II occupancy (assessed by ChIP-seq) versus fold change (teriflunomide/DMSO) in mRNA abundance (assessed by RNAseq) following 48-hour treatment of A375 cells with teriflunomide or DMSO vehicle control; data derived from ^38^.

One potential target that met the selection criteria was CDK9, which together with cyclin T1 or T2 forms positive transcription elongation factor B (P-TEFb). P-TEFb is required for the release of promoter-proximal paused RNA polymerase II (Pol II) into productive elongation and therefore is essential for Pol II transcription from paused promoters ^48,49^. The potent P-TEFb inhibitor flavopiridol ^50^ phenocopied BMS-345541 in our assays, as it blocked APP induction downstream of DHODH, IMPDH1/2 (by mycophenolate), or CTP synthase (by 3-deazauridine ^51^) inhibition (Fig S4A). This suggests that APP induction downstream of nucleotide starvation requires P-TEFb-mediated paused Pol II release. It also suggests that the BMS-345541 effect of reversing BQ-induced APP upregulation is due to P-TEFb inhibition.

Within the list of kinases bound by BMS-345541 (Fig 4A) we eliminated those that were a) not expressed by CFPAC-1 cells in our RNA-seq data, b) not bound by flavopiridol in KINOMEscan data, or c) bound by ruxolitinib in KINOMEscan data with Kd < 500 nM (as 2.5 μM ruxolitinib failed to reverse BQ-mediated APP induction (Fig 3A)). Five candidates (besides CDK9) remained that were bound by both BMS-345541 and flavopiridol in KINOMEscan assays. Of these, three are CDKs known to play a role in transcription (CDK7, CDK13, and CDK16). Inhibition of any of these CDKs could theoretically account for the observed effects of flavopiridol and BMS-345541. However, previous studies suggest that flavopiridol inhibition of these CDKs in vivo is much less efficient than in cell-free assays because it is competitive with ATP (and thus less efficient in living cells where the ATP concentration is in the 1-10 mM range, which is much higher than in cell-free assay conditions), while its inhibition of P-TEFb is not affected by ATP concentration ^50^. Furthermore, flavopiridol and the CDK7 inhibitor THZ1 have very different (and mutually exclusive) effects on transcriptional processes ^52^, arguing against CDK7 inhibition as the mechanism of flavopiridol’s effect.

To further probe whether the observed effect of flavopiridol was due to CDK9 inhibition, we tested two other CDK9 inhibitors (AT7519 and dinaciclib). Both CDK9 inhibitors phenocopied flavopiridol in our assays (Fig 4B). Furthermore, a previously characterized CDK9-targeted proteolysis targeting chimera (PROTAC), termed PROTAC2 ^53^, had the same effect (Fig 4B). PROTAC2 consists of a CDK9-binding aminopyrazole warhead conjugated to pomalidomide, which recruits the E3 ubiquitin ligase Cereblon (*CRBN*). Cereblon in turn ubiquitinates CDK9, resulting in its proteasomal degradation. Co-treatment of HEK-293 cells with PROTAC2 and pomalidomide prevents PROTAC2-mediated CDK9 degradation, as free pomalidomide competes with PROTAC2 for Cereblon binding ^53^. We observed that PROTAC2 (1 μM) blocked BQ-mediated APP induction, and this effect was reversed by co-treatment with 10-fold excess pomalidomide (10 μM); however, when we increased the concentration of PROTAC2 to 10 μM (so that PROTAC2 and pomalidomide concentrations were equal), pomalidomide no longer had this effect (Fig 4C). Consistently, immunoblot analysis showed that 10 μM pomalidomide prevents CDK9 degradation upon 1 μM but not 10 μM PROTAC2 treatment (Fig 4D). When we repeated the experiment shown in Fig 4B with HCT116 cells, we found that all CKD9 inhibitors reversed BQ-mediated APP induction, but PROTAC2 did not (Fig 4E). Concordantly, immunoblot analysis showed that PROTAC2 did not cause CDK9 depletion in HCT116 cells treated in parallel (Fig 4F).

Taken together, these results demonstrate that CDK9 degradation is necessary for the reversal effect of PROTAC2 and that CDK9 is required for BQ-mediated APP induction.

The dependence of BQ-mediated APP induction on CDK9 strongly suggests that nucleotide starvation enforces nascent transcription of APP genes, as opposed to increased mRNA stability. This is further supported by the rapid buildup of APP transcripts following DHODH inhibitor treatment (within 4 hours, Fig S3A). Additionally, ChIP-seq analysis of global Pol II occupancy following 48-hour teriflunomide treatment in A375 cells ^38^ shows increased Pol II occupancy across many APP genes, and fold change in Pol II occupancy significantly correlated with fold change in mRNA expression under the same conditions (Fig 4G). Overall, these results show that nucleotide starvation induces an antigen presentation gene expression program that is independent of canonical APP regulators but depends on CDK9/P-TEFb.

### BQ suppresses tumor growth, induces MHC-I expression, and increases immunotherapy efficacy in a syngeneic melanoma model

Enforced MHC-I upregulation by various interventions can facilitate anticancer immunity and enhance the efficacy of immune checkpoint blockade (ICB) by antibodies directed at PD-(L)1 and/or CTLA-4 ^24-27^. Moreover, high MHC-I expression has been proposed as a predictor of ICB response ^28-31^, and high expression of MHC-I and other APP genes, including *NLRC5* and *TAP1*, correlates with better survival in patients with melanoma (Fig S5A), for whom ICB is a first-line therapy. Therefore, we asked if BQ could improve anticancer immunity in the B16F10 melanoma immunocompetent mouse model, which is typically refractory to dual ICB (i.e., anti-PD-1 plus anti-CTLA-4) ^54^.

BQ (10 mg/kg daily IP injection) markedly suppressed tumor growth and led to reduced tumor burden (Fig 5A-B). Historically, the lead tool compound that was ultimately modified to BQ (called NSC 339768) was prioritized in part based on its activity against B16 melanoma ^55^; however, to our knowledge, this is the first direct demonstration of BQ activity in this model system. Consistent with our *in vitro* metabolomics data (Fig 1I-J, S1B), BQ treatment caused marked buildup of metabolites upstream of DHODH and depletion of downstream pyrimidine nucleotide species in B16F10 tumors (Fig 5C), confirming target engagement *in vivo*. Metabolomics analysis of BQ- and vehicle-treated tumors separated in principal component analysis (Fig S5B) and unsupervised hierarchical clustering (Fig S5C), confirming the perturbation of tumor metabolism following DHODH inhibition.

**Figure 5:**
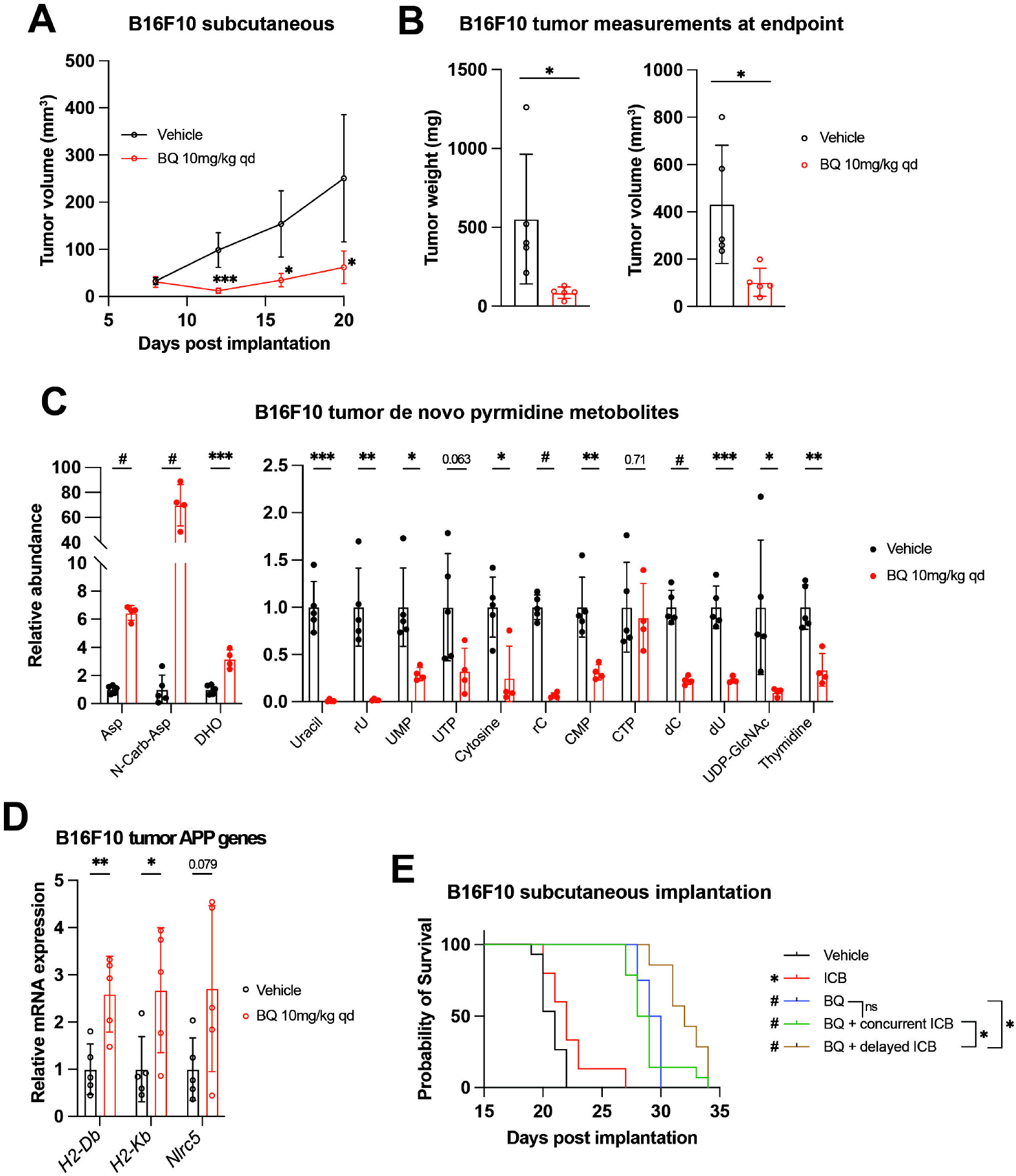
BQ inhibits tumor growth, increases tumor MHC-I, and enhances immune checkpoint blockade efficacy in B16F10 murine melanoma model. **A-D)** B16F10 cells were injected subcutaneously into syngeneic hosts and mice were treated with brequinar (BQ; 10mg/kg IP daily) or vehicle control starting at day 7 post implantation. **A)** Longitudinal measurement of B16F10 subcutaneous tumors with BQ (10mg/kg IP daily) or vehicle treatment. **B)** Weight (left) and volume (right) of tumors at necropsy. **C)** Quantification of metabolites from B16F10 tumors harvested at necropsy. **D)** RT-qPCR analysis of APP genes from tumors harvested at necropsy. For (A-D), data represent mean +/-SD of n = 5 mice per group (n = 4 for BQ group in C). * indicates p < 0.05, ** indicates p < 0.01, *** indicates p < 0.001, and # indicates p < 0.0001 by unpaired t-test. **E)** Kaplan-Meier survival analysis for mice implanted with B16F10 tumors as in (A-D) and treated with indicated regimens; see Fig S5D for treatment timeline. * indicates p < 0.05, # p < 0.0001 by Mantel-Cox logrank test. For vehicle, immune checkpoint blockade (ICB; Anti-CTLA-4 and anti-PD-1; 100μg/mouse each, IP twice per week), and BQ (10mg/kg IP daily) + concurrent ICB, n = 15. For BQ, n = 7. For BQ + delayed ICB, n = 8.

BQ-treated B16F10 tumors showed increased mRNA expression of MHC-I (*H2-Db* and *H2-Kb*) and *Nlrc5* (Fig 5D). We thus addressed whether BQ could augment the efficacy of dual ICB (anti-CTLA-4 plus anti-PD-1) with the knowledge that enforced MHC-I antigen presentation has also been shown to boost the effect of ICB ^24,26,27^. While BQ is not an approved medication, two FDA-approved low potency DHODH inhibitors (leflunomide, teriflunomide) are effective in treating autoimmune conditions such as rheumatoid arthritis and multiple sclerosis and act to decrease the activity of auto-reactive T-lymphocytes ^56-58^. It was possible that BQ treatment may actually impair the effectiveness of ICB by inhibiting T-lymphocytes despite augmented cancer cell antigen presentation. We, therefore, tested both concurrent, upfront administration of BQ plus dual ICB and sequential administration of BQ followed by dual ICB (Fig S5D).

Similar to its impressive activity in our first experiment (Fig 5A-B), BQ monotherapy conferred marked survival benefit. This was significantly enhanced by subsequent dual ICB, while dual ICB alone conferred only marginally prolonged survival, and concurrent BQ plus dual ICB did not significantly improve survival versus BQ monotherapy (Fig 5E). This suggests that sequential (rather than concurrent) administration of DHODH inhibitor and ICB may be superior. Hypotheses that may explain these findings include: a) Concurrent BQ dampens the initial anticancer immune response generated by dual ICB, or b) cancer cell MHC-I and related genes are not maximally upregulated at the time of ICB administration with concurrent treatment. Taken together, these results show that BQ causes pyrimidine nucleotide depletion, MHC-I and APP gene transcriptional upregulation, and additive survival benefit with dual ICB in a highly aggressive and ICB-refractory mouse melanoma model.

## Discussion

Our results demonstrate that pyrimidine nucleotide depletion by DHODH inhibition causes increased expression of APP genes and increased antigen presentation via MHC-I across a diverse panel of cancer cell lines (Fig 1). This effect of BQ and teriflunomide is strictly dependent on pyrimidine nucleotide depletion, as it was abrogated by restoration of pyrimidine levels with exogenous uridine (Fig 2B-D and Fig S2B-E). Furthermore, genetic deletion of *DHODH* recapitulated this effect, and treatment of DHODH knockout cells with BQ did not further increase MHC-I mRNA expression (Fig 2E). Our inhibitor reversal studies determined that BQ-mediated APP induction is independent of several canonical APP regulatory pathways, including IFN-JAK-STAT, cGAS-STING-TBK1, and NF-ĸB (Fig 3 and Fig S3). We showed that this effect relies on P-TEFb-mediated release of Pol II from promoter-proximal paused state to productive elongation (Fig 4). These findings were extended to inhibition of IMPDH (which depletes cellular GTP) and CTPS (which depletes cellular CTP), as these effects were also reversible with P-TEFb inhibition (Fig S4A). This suggests that pharmacologic depletion of these nucleotide species also triggers APP upregulation in a P-TEFb-dependent manner.

Since T-cell recognition of antigens via MHC-I is required for T-cell-mediated elimination of cancer cells or virus-infected cells, these results have important implications for the development of nucleotide synthesis inhibitors as anticancer/antiviral therapies. We provide proof of concept evidence that pretreatment with DHODH inhibitors can improve the efficacy of immune checkpoint blockade in a highly aggressive and ICB-refractory mouse melanoma model (Fig 5 and Fig S5). Because BQ-mediated APP induction does not require interferon signaling, this strategy may have particular relevance for clinical scenarios in which tumor antigen presentation is dampened by the loss or silencing of cancer cell interferon signaling, which has been demonstrated to confer both intrinsic ^59^ and acquired ^23^ ICB resistance in human melanoma patients.

Emerging evidence suggests that cancer cell MHC-I expression predicts favorable response to ICB, and several recent studies have shown that enforced cancer cell MHC-I expression enhances anticancer immunity and ICB efficacy in various mouse models. Accordingly, functional genomic screens for regulators of cancer cell MHC-I expression have recently been undertaken, and these efforts have revealed novel molecular targets to induce cancer cell APP activity ^27,60^. Agents shown to increase cancer cell antigen presentation include hydroxychloroquine (by autophagy inhibition) ^24^, poly(I:C) (by NF-ĸB activation downstream of dsRNA sensing) ^26^, SMAC mimetics (by NF-ĸB activation) ^27^, CDK4/6 inhibitors (by activation of endogenous genomic retroviral elements) ^25^, and others. It is very likely that many other anticancer drugs perturb cancer cell antigen presentation and/or have other immunomodulatory properties in addition to their cell-intrinsic antiproliferative activity ^61^, and this area requires further scrutiny. In this study, we identified DHODH inhibition as a powerful inducer of antigen presentation and MHC-I expression in diverse cancer cell lines and in HEK-293T cells.

Previous studies have linked pyrimidine depletion with upregulation of innate immunity and interferon-stimulated genes ^33,34^, and this was confirmed by our transcriptomic profiling experiments (Fig 1B-C). Multiple mechanistic explanations for these observations have been suggested. Lucas-Hourani et al. proposed that interferon-stimulated gene expression requires the DNA damage checkpoint kinase ATM ^33^, while Sprenger et al. conclude that pyrimidine depletion causes accumulation of mitochondrial DNA in the cytosol, which is sensed by the cGAS-STING-TBK1 pathway to promote innate immunity ^34^. In our models, neither ATM/ATR nor TBK1 inhibition blocked BQ-mediated APP induction (Fig 4A). It is possible that pyrimidine nucleotide shortage leads to APP induction by multiple redundant mechanisms, any of which may predominate based on the cellular context. We speculate that cells may have evolved multiple means of sensing acute pyrimidine shortage as a way to detect viral infection or malignant transformation, as both viral replication and uncontrolled cell proliferation avidly consume nucleotides.

Our focused chemical screen for MHC-I inducers (Fig 2F) identified the approved IMPDH1/2 inhibitor mycophenolate, which was subsequently validated in multiple other cell types (Fig S2D, S4A). This is consistent with a recent study in which IMPDH inhibition was shown to enhance ICB efficacy by favorably altering the MHC-I peptide repertoire and increasing immunoproteasome expression ^62^. However, in this study, the cancer cells were pretreated with IMPDH inhibitor before implantation into syngeneic hosts, and so possible countervailing immunosuppression by systemic IMPDH inhibitor treatment was not addressed ^62^. Our *in vivo* results (Fig 5F) highlight the importance of timing/sequence when administering immunotherapy in combination with nucleotide synthesis inhibitors and suggest that upfront BQ followed by ICB may be superior to concurrent administration.

Thymidylate synthase inhibition was recently shown to induce MHC-I in a model of diffuse large B cell lymphoma ^60^. The failure of thymidylate synthase inhibitors 5-fluorouracil and methotrexate to induce MHC-I in our screen (Fig 2F) may be attributable to cell line differences (PDAC vs DLBCL), dose/duration considerations, or the use of different thymidylate synthase inhibitors than in their study (which used pemetrexed and raltitrexed). Thus, it appears that the abundance of multiple nucleotide species can exert context-dependent influence on MHC and APP gene expression, and key details of this relationship remain to be elucidated.

Overall, our study establishes P-TEFb and Pol II elongation control as a mechanistic link between nucleotide depletion and APP induction. We provide proof of concept evidence for combinatorial benefit of DHODH inhibition and immune checkpoint blockade in an aggressive and poorly immunogenic mouse model of melanoma. A deeper understanding of metabolic control of antigen presentation will enable rational therapy development for cancer and viral infection.

## Materials and Methods

### Cell culture and cell lines

The S2-013 cell line is a clonal derivative of the Suit2 cell line and was a kind gift from the Tony Hollingsworth laboratory at the University of Nebraska Medical Center. The MiaPaCa2 *IKK2*-KO and parental wild-type MiaPaCa2 cell lines were a kind gift from the Amar Natarajan laboratory at the University of Nebraska Medical Center. All other cell lines in this study were obtained from American Type Culture Collection (Manassas, VA, USA). All human cell lines were authenticated by STR profiling by the Genetics Core at the University of Arizona. Cells were routinely (at the time of initial revival from liquid nitrogen storage and at least every 6 months) determined to be free of mycoplasma contamination by PCR-based methods. Cells were cultured in Dulbecco’s modified Eagle medium (Sigma-Aldrich, St Louis, MO, USA) supplemented with 50 IU/mL penicillin, 50[μg/mL streptomycin, and incubated at 37[°C in a humidified incubator with 5% CO_2_. Cells were maintained at 10% fetal bovine serum (FBS). Upon reaching 70–80% confluency, cells were passaged by washing with phosphate-buffered saline (PBS) before adding 0.25% trypsin (Caisson Labs, Smithfield, UT, USA) and plating at 25% confluency.

### Drug treatment of cultured cells for RT-qPCR and flow cytometry experiments

Drug treatment dose and duration are indicated for each experiment. A table of manufacturer and catalog number for each agent described can be found in Supplementary Table 1. For stimulation with poly(dA:dT), 2μg of poly(dA:dT) and 2μl of Lipofectamine2000 (Invitrogen #11668027) were incubated in 400μl Opti-MEM (Gibco #11058021) for 30 minutes at room temperature and then added to cells in 2ml final volume of complete media.

### Cell viability assays

Cells were seeded in 96 well plates (1000 cells per well in 90μl media) and allowed to equilibrate overnight. Cells were then treated with indicated compounds (final volume 100μl) for 72 hours, and viability was assessed by CellTiter-Glo assay (Promega, Madison, WI). Luminescence values for each condition were normalized to the average luminescence of the vehicle-treated control replicates. Experiments were performed at least three times using biological triplicates for each condition. Dose-response curves were fit to nonlinear regression models using Prism9 software.

### Liquid chromatography – tandem mass spectrometry-based metabolomics analysis

For in vitro metabolomics experiments, 5 x 10^5^ cells were seeded in 6-well plates and allowed to equilibrate overnight. At the start of each assay, the cell culture media was changed, and fresh media with desired conditions was added (to eliminate metabolite depletion from overnight equilibration as a confounding variable). Following 8-hour treatment of cancer cell lines with BQ (or in the case of Fig 2B, 24-hour treatment with BQ +/-100μM uridine), polar metabolites were extracted and quantified as previously described ^63^. For B16F10 tumor metabolomics, subcutaneous tumors were harvested at necropsy and immediately snap frozen in liquid nitrogen and stored at -80 °C. Tumors were subsequently ground into fine powder in liquid nitrogen using a mortar and pestle, and metabolites were extracted using the same method as for cultured cells. Peak areas were normalized to the mass of tumor tissue that was input.

Datasets were processed using Skyline (MacCoss Lab Software), and Metaboanalyst5.0 web tool was used to generate principal component analysis and heatmap visualizations of resulting datasets. Raw LC-MS/MS data and R-command history for Metaboanalyst analysis are available upon request. Relative metabolite abundances were normalized to the average peak area of the experimental control group and were compared using two-way ANOVA with Bonferonni’s post-test correction for multiple comparisons. P < 0.05 was considered significant.

### Mice studies

All procedures were approved by the Institutional Animal Care and Use Committee (IACUC) at the University of Nebraska Medical Center. For tumor xenograft studies, 10^4^ B16F10 cells in a 1:1 vol/vol ratio (100μl final volume) with Matrigel were injected subcutaneously into the right flank of 10-week-old female C57BL/6J mice (Jackson labs). Tumors of live mice were serially measured in two dimensions using digital calipers, and tumor volume for Fig 5A was calculated as (0.5L x W^2^), where L is the long dimension and W is the short dimension. For Fig 5B-C, tumors were harvested at necropsy, weighed on an analytical balance (for Fig 5B), and measured in three perpendicular dimensions by calipers to generate volume measurements for Fig 5C, which were calculated as (dimension 1 x dimension 2 x dimension 3).

For survival experiments (Fig 5F), mice were monitored daily for signs of euthanasia criteria or actual demise. When tumor volume reached 2000cm^3^ as determined by the above formula for live mice (0.5L x W^2^), mice were sacrificed according to protocol euthanasia criteria.

Brequinar was obtained from Clear Creek Bio and dissolved in 0.9% NaCl. For both endpoint and survival studies, BQ (10mg/kg) or vehicle solvent (0.9% NaCl) was injected intraperitoneally daily. Anti-CTLA-4 and anti-PD-1 antibodies, as well as their respective isotype controls, were obtained from BioXCell. Antibodies were dosed at 100μg/mouse IP twice per week.

### RNA sequencing and gene set enrichment analysis

For RNA sequencing experiments, S2-013 or CFPAC-1 cells were treated with brequinar for the indicated dose and duration (Fig 1 and S1). For two-week drug treatment experiments, cells were passaged every 3 days and 5 x 10^5^ cells were reseeded in a new 10cm tissue culture dish. RNA was isolated using RNEasy Mini kit (Qiagen, catalog number 74104).

Samples were processed by BGI Genomics (San Jose, California, USA) according to their proprietary method. Briefly, RNA quality check was performed using Agilent 2100 Bioanalyzer. Poly-A-containing mRNA was isolated using magnetic beads and then fragmented using divalent cations under elevated temperature. cDNA synthesis was performed using reverse transcriptase and RNase H. Adapter sequences were then ligated onto cDNA fragments, purified, enriched by PCR, quantified by Qubit, and pooled to generate the final library. Libraries were then sequenced using the BGI DNBseq platform. Reads mapped to rRNA, low quality reads, and reads with adaptors were removed. The resulting clean reads were mapped to the reference genome (hg19_UCSC_20180115) using HISAT2 program (http://www.ccb.jhu.edu/software/hisat/index.shtml) and converted to fragments for kilobase per million mapped reads (FPKM).

Fold change FPKM (brequinar/vehicle control) values for all expressed genes (FPKM > 1) were subjected to gene set enrichment analysis ^36^ with GSEA prerank using HALLMARK and KEGG genes sets from the Molecular Signatures Database (MSigDB) as previously described ^64^. Gene sets positive enriched with FDR q < 0.25 are shown in Fig 1B.

### Real time quantitative PCR analysis for mRNA expression

For in vitro RT-qPCR experiments, RNA was harvested using Trizol reagent (Thermo Fisher Scientific, Waltham, MA, USA) according to manufacturer’s instructions. For tumor RT-qPCR, tumors were crushed with mortar and pestle in liquid nitrogen, and Trizol was used to extract RNA from the resulting powder, just as for cultured cells. cDNA synthesis was performed (1μg RNA input) using BioRad (Hercules, CA, USA) iScript cDNA synthesis kit (catalog number 1708891) according to manufacturer’s instructions. For RT-qPCR reactions, 3μl of diluted cDNA, 2μl of primer mix (diluted to a final concentration of 200 nM for forward and reverse primers), and 5μl SYBR green master mix (cat #) were mixed (10μl final volume), and reactions were analyzed using Applied Biosystems QuantStudio5 instrument with previously reported thermocycling parameters ^65^.

18S rRNA was used as a loading control to generate delta Ct values, and each sample was normalized to the experimental control delta Ct values to generate delta delta Ct values which were converted to fold change by (2^^^-ddCt). For all experiments, *ACTB* (beta-actin) mRNA expression was quantified and used as an additional loading control, and results were concordant regardless of whether 18S or *ACTB* was used for normalization. For Fig 5E, each data point represents the average expression value of 4 technical replicates for an individual tumor.

For pairwise comparisons, an unpaired student’s t-test was used. For comparisons of 3 or more conditions, a two-way ANOVA with Bonferonni’s post-test correction for multiple comparisons was used. P < 0.05 was considered significant. Primer sequences for RT-qPCR reactions are provided in Supplementary Table 2.

### Flow cytometry measurement of cell surface MHC-I

Cells were treated as described and then detached with Accutase (Sigma Aldrich #A6964), washed twice with PBS, stained with fluorescent dye-conjugated antibodies against H2-Db (BioLegend #111508) or HLA-A/B/C (BioLegend #311418, BioLegend #311406) for 30 minutes at 4 °C in PBS, washed once more with PBS, and then resuspended in FACS buffer and subjected to flow cytometry analysis for fluorescence intensity. Aqua live/dead dye (Invitrogen #L34957) or propidium iodide was used to exclude dead cells from the analysis.

### Western blot

Protein isolation from cultured cells and western blotting procedure were described previously ^63^. CDK9 antibody was obtained from Cell Signaling Technology (catalog number 2316, clone C12F7), HSP70 antibody was obtained from Cell Signaling Technology (catalog number 4872), and beta-actin antibody was obtained from Santa Cruz Biotechnology (catalog number sc-4778, clone C4). Blots were incubated with primary antibody overnight at 4 °C, washed, incubated with secondary antibody conjugated with horseradish peroxidase for 45 min at room temperature, washed, developed with ECL reagent and visualized by autoradiography using plain film.

### Procurement and analysis of previously published datasets

All datasets reported by Tan and colleagues ^38^ were obtained from Gene Expression Omnibus, accession numbers GSE68053 and GSE68039. Processed RNA sequencing data for human A375 melanoma cells treated with DMSO vehicle control (GSM1661518, GSM1661518), or teriflunomide (25μM) for 12 hours (GSM1661510, GSM1661511), 24 hours (GSM1661512, GSM1661513), 48 hours (GSM1661514, GSM1661515), or 72 hours (GSM1661516, GSM1661517) was downloaded as an Excel file from GSE68039 (GSE6809_A375.FPKM.xls) and directly analyzed by manual inspection. The two FPKM values for each experimental condition were averaged, and these average values were used to calculate the fold change (teriflunomide/DMSO) values presented in Figure 1E and Figure 4G.

For chromatin immunoprecitation sequencing (ChIP-seq) datasets (used to generate Fig 4G), Fastq files for human A375 melanoma cells treated for 48 hours with DMSO (GSM1661790) or teriflunomide (GSM1661791) for were downloaded from GSE68053, trimmed of adapter sequences at the 3′ends with trim_galore v0.6 (https://github.com/FelixKrueger/TrimGalore), and aligned to hg38 using Bowtie (v1.2.3) (ref = https://genomebiology.biomedcentral.com/articles/10.1186/gb-2009-10-3-r25) with parameters --minins 18 --maxins 1000 --fr --best --allow-contain. Reads overlapping with the longest transcript of each gene (Genecode v32 and https://github.com/GeoffSCollins/PolTools/blob/master/PolTools/static/longest_transcript_with_downstream_start_codon.txt) were counted with BEDtools intersect (v2.27.1). Library size correction factors were calculated separately for the ChIP-seq datasets. The correction factor for a given ChIP-seq sample was computed by dividing the number of mapped reads in that sample by the average number of mapped reads across all ChIP samples (DMSO, A771726). After normalization, the total number of read counts (now corrected for total number of mapped reads per sample) aligned to each gene of interest were used to calculate fold change (teriflunomide/DMSO) in Pol II occupancy values presented in Figure 4G.

## ACKNOWLEDGMENTS

This work was supported in part by funding from the National Institutes of Health, including R01CA163649 and U54CA274329 to PKS, F30CA265277 to NJM, R21CA251151 to AN, and R35GM126908 to DHP.

## Figure legends

**Figure S1: BQ treatment upregulates APP genes and depletes pyrimidine nucleotides. A)** Heatmap showing log2 fold change mRNA expression of APP genes in S2-013 cells treated with BQ for indicated dose and duration. **B-C)** RT-qPCR quantification of APP genes after two-week BQ treatment of CFPAC-1 (250nM) (B) or B16F10 (10μM) (C) cells. **D-E)** Quantification of pyrimidine metabolites in CFPAC-1 (C) or B16F10 (D) cells treated with BQ for 8 hours at indicated doses. Data represent mean +/-SEM of four (CFPAC-1) or six (B16F10) biological replicates. * indicates p < 0.05, ** p < 0.01, *** p < 0.001, and # p < 0.0001 by unpaired t-test (B, C) or two-way ANOVA with Bonferonni’s post-comparison test (D, E).

**Figure S2: Uridine rescues B16F10 cells from teriflunomide toxicity but does not alter APP expression by itself. A)** Dose-response cell viability experiment as in Fig 2A but with teriflunomide (Ter) instead of BQ. **B)** RT-qPCR analysis of indicated genes following treatment with teriflunomide +/-uridine (1mM) for 24 hours. Data represent mean +/-SEM of three determinations. One representative result of three independent experiments is shown. **C)** RT-qPCR analysis of indicated genes after treatment with vehicle or uridine (1mM) for 24 hours. Data represent mean +/-SEM of four determinations. One representative result of three independent experiments is shown. **D)** RT-qPCR analysis of indicated genes following treatment with BQ (10μM) +/-uridine (1mM) for 24 hours. Data represent mean +/-SD of three independent experiments. **E)** Flow cytometry analysis of cell surface MHC-I (H2-Db) following 24-hour uridine (1mM) treatment. Data represent mean +/-SEM of three independent experiments. **F)** RT-qPCR analysis of indicated genes in B16F10 cells following 24-hour treatment with MPA (5μM). Data represent mean +/-SEM of four determinations. One representative result of three independent experiments is shown.

**Figure S3: A)** RT-qPCR time course analysis for MHC-I genes in HEK-293T cells treated for indicated times with BQ (10μM). Data represent mean +/-SD of four determinations. **B)** RT-qPCR analysis for *Nlrc5* (left) or *Tap1* (right) in B16F10 cells treated for 24 hours with indicated agents. Data represent mean +/-SD of four determinations. # indicates p < 0.0001 with two-way ANOVA with Bonferroni post-comparison test. Representative results for one of three independent experiments are shown. **C)** Heatmap indicating RT-qPCR analysis for indicated genes in wild-type or *IKK2*-KO MiaPaCa2 cells treated with indicated agents for 24 hours. Numbers in the heatmap represent mean of four determinations.

**Figure S4: A)** RT-qPCR analysis for indicated genes in HCT116 cells treated with indicated agents in the presence or absence of flavopiridol (1μM). Numbers in the heatmap represent mean of three determinations.

**Figure S5: A)** Overall survival in melanoma patients (SKCM) from the Cancer Genome Atlas (TCGA) with above (indicated in red) and below median (indicated in blue) mRNA expression of indicated genes. **B-C)** Principal component analysis (PCA) plot (B) and unsupervised hierarchical clustering (C) from metabolomics analysis of B16F10 tumors at necropsy (Fig 5C). **D)** Treatment regimen of mice from Fig 5E.

## References

1 Warburg, O. On the origin of cancer cells. Science 123, 309–314, doi:10.1126/science.123.3191.309 (1956).

2 Hanahan, D. & Weinberg Robert A. Hallmarks of Cancer: The Next Generation. Cell 144, 646–74, 10.1016/j.cell.2011.02.013(2011).

3 Mullen, N. J. & Singh, P. K. Nucleotide metabolism: a pan-cancer metabolic dependency. Nature Reviews Cancer 23, 275–294, doi:10.1038/s41568-023-00557-7 (2023).

4 Wang, W., Cui, J., Ma, H., Lu, W. & Huang, J. Targeting Pyrimidine Metabolism in the Era of Precision Cancer Medicine. Frontiers in Oncology 11, doi:10.3389/fonc.2021.684961 (2021).

5 Shukla, S. K. et al. MUC1 and HIF-1alpha Signaling Crosstalk Induces Anabolic Glucose Metabolism to Impart Gemcitabine Resistance to Pancreatic Cancer. Cancer Cell 32, 71–87.e77, doi:10.1016/j.ccell.2017.06.004(2017.

6 Christian, S. et al. The novel dihydroorotate dehydrogenase (DHODH) inhibitor BAY 2402234 triggers differentiation and is effective in the treatment of myeloid malignancies. Leukemia 33, 2403–2415, doi:10.1038/s41375-019-0461-5 (2019).

7 Sykes, D. B. et al. Inhibition of Dihydroorotate Dehydrogenase Overcomes Differentiation Blockade in Acute Myeloid Leukemia. Cell 167, 171–186.e115, doi:10.1016/j.cell.2016.08.057 (2016).

8 Wang, X. et al. Targeting pyrimidine synthesis accentuates molecular therapy response in glioblastoma stem cells. Science Translational Medicine 11, eaau4972, doi:10.1126/scitranslmed.aau4972 (2019).

9 Koundinya, M. et al. Dependence on the Pyrimidine Biosynthetic Enzyme DHODH Is a Synthetic Lethal Vulnerability in Mutant <em>KRAS</em>-Driven Cancers. Cell Chemical Biology 25, 705–717.e711, doi:10.1016/j.chembiol.2018.03.005 (2018).

10 Santana-Codina, N. et al. Oncogenic KRAS supports pancreatic cancer through regulation of nucleotide synthesis. Nat Commun 9, 4945, doi:10.1038/s41467-018-07472-8 (2018).

11 Brown, K. K., Spinelli, J. B., Asara, J. M. & Toker, A. Adaptive Reprogramming of <em>De Novo</em> Pyrimidine Synthesis Is a Metabolic Vulnerability in Triple-Negative Breast Cancer. Cancer Discovery 7, 391–399, doi:10.1158/2159-8290.Cd-16-0611 (2017).

12 Mathur, D. et al. PTEN Regulates Glutamine Flux to Pyrimidine Synthesis and Sensitivity to Dihydroorotate Dehydrogenase Inhibition. Cancer Discovery 7, 380–390, doi:10.1158/2159-8290.Cd-16-0612 (2017).

13 Li, L. et al. Identification of DHODH as a therapeutic target in small cell lung cancer. Science Translational Medicine 11, eaaw7852, doi:10.1126/scitranslmed.aaw7852 (2019).

14 Bajzikova, M. et al. Reactivation of Dihydroorotate Dehydrogenase-Driven Pyrimidine Biosynthesis Restores Tumor Growth of Respiration-Deficient Cancer Cells. Cell Metab 29, 399–416.e310, doi:10.1016/j.cmet.2018.10.014 (2019).

15 Falzone, L., Salomone, S. & Libra, M. Evolution of Cancer Pharmacological Treatments at the Turn of the Third Millennium. Frontiers in Pharmacology 9, doi:10.3389/fphar.2018.01300 (2018).

16 Vaddepally, R. K., Kharel, P., Pandey, R., Garje, R. & Chandra, A. B. Review of Indications of FDA-Approved Immune Checkpoint Inhibitors per NCCN Guidelines with the Level of Evidence. Cancers (Basel) 12, doi:10.3390/cancers12030738 (2020).

17 Mundry, C. S., Eberle, K. C., Singh, P. K., Hollingsworth, M. A. & Mehla, K. Local and systemic immunosuppression in pancreatic cancer: Targeting the stalwarts in tumor’s arsenal. Biochim Biophys Acta Rev Cancer 1874, 188387, doi:10.1016/j.bbcan.2020.188387 (2020).

18 Waldman, A. D., Fritz, J. M. & Lenardo, M. J. A guide to cancer immunotherapy: from T cell basic science to clinical practice. Nat Rev Immunol 20, 651–668, doi:10.1038/s41577-020-0306-5 (2020).

19 Pishesha, N., Harmand, T. J. & Ploegh, H. L. A guide to antigen processing and presentation. Nature Reviews Immunology 22, 751–764, doi:10.1038/s41577-022-00707-2 (2022).

20 Cornel, A. M., Mimpen, I. L. & Nierkens, S. MHC Class I Downregulation in Cancer: Underlying Mechanisms and Potential Targets for Cancer Immunotherapy. Cancers (Basel) 12, doi:10.3390/cancers12071760 (2020).

21 Dhatchinamoorthy, K., Colbert, J. D. & Rock, K. L. Cancer Immune Evasion Through Loss of MHC Class I Antigen Presentation. Front Immunol 12, 636568, doi:10.3389/fimmu.2021.636568 (2021).

22 Han, P. et al. Genome-Wide CRISPR Screening Identifies JAK1 Deficiency as a Mechanism of T-Cell Resistance. Front Immunol 10, 251, doi:10.3389/fimmu.2019.00251 (2019).

23 Zaretsky, J. M. et al. Mutations Associated with Acquired Resistance to PD-1 Blockade in Melanoma. N Engl J Med 375, 819–829, doi:10.1056/NEJMoa1604958 (2016).

24 Yamamoto, K. et al. Autophagy promotes immune evasion of pancreatic cancer by degrading MHC-I. Nature 581, 100–105, doi:10.1038/s41586-020-2229-5 (2020).

25 Goel, S. et al. CDK4/6 inhibition triggers anti-tumour immunity. Nature 548, 471–475, doi:10.1038/nature23465 (2017).

26 Kalbasi, A. et al. Uncoupling interferon signaling and antigen presentation to overcome immunotherapy resistance due to JAK1 loss in melanoma. Sci Transl Med 12, doi:10.1126/scitranslmed.abb0152 (2020).

27 Gu, S. S. et al. Therapeutically Increasing MHC-I Expression Potentiates Immune Checkpoint Blockade. Cancer Discov 11, 1524–1541, doi:10.1158/2159-8290.Cd-20-0812 (2021).

28 Rodig, S. J. et al. MHC proteins confer differential sensitivity to CTLA-4 and PD-1 blockade in untreated metastatic melanoma. Sci Transl Med 10, doi:10.1126/scitranslmed.aar3342 (2018).

29 Liu, D. et al. Integrative molecular and clinical modeling of clinical outcomes to PD1 blockade in patients with metastatic melanoma. Nature Medicine 25, 1916–1927, doi:10.1038/s41591-019-0654-5 (2019).

30 Grasso, C. S. et al. Conserved Interferon-γ Signaling Drives Clinical Response to Immune Checkpoint Blockade Therapy in Melanoma. Cancer Cell 38, 500–515.e503, doi:10.1016/j.ccell.2020.08.005 (2020).

31 Shklovskaya, E. et al. Tumor MHC Expression Guides First-Line Immunotherapy Selection in Melanoma. Cancers (Basel) 12, doi:10.3390/cancers12113374 (2020).

32 Luthra, P. et al. Inhibiting pyrimidine biosynthesis impairs Ebola virus replication through depletion of nucleoside pools and activation of innate immune responses. Antiviral Res 158, 288–302, doi:10.1016/j.antiviral.2018.08.012 (2018).

33 Lucas-Hourani, M. et al. Inhibition of Pyrimidine Biosynthesis Pathway Suppresses Viral Growth through Innate Immunity. PLOS Pathogens 9, e1003678, doi:10.1371/journal.ppat.1003678 (2013).

34 Sprenger, H.-G. et al. Cellular pyrimidine imbalance triggers mitochondrial DNA–dependent innate immunity. Nature Metabolism 3, 636–650, doi:10.1038/s42255-021-00385-9 (2021).

35 Liberzon, A. et al. Molecular signatures database (MSigDB) 3.0. Bioinformatics 27, 1739–1740, doi:10.1093/bioinformatics/btr260 (2011).

36 Subramanian, A. et al. Gene set enrichment analysis: A knowledge-based approach for interpreting genome-wide expression profiles. Proceedings of the National Academy of Sciences 102, 15545–15550, doi:doi:10.1073/pnas.0506580102 (2005).

37 Mullen, N. J. et al. ENT1 blockade by CNX-774 overcomes resistance to DHODH inhibition in pancreatic cancer. Cancer Letters 552, 215981, 10.1016/j.canlet.2022.215981(2023).

38 Tan, Justin L. et al. Stress from Nucleotide Depletion Activates the Transcriptional Regulator HEXIM1 to Suppress Melanoma. Molecular Cell 62, 34–46, doi:10.1016/j.molcel.2016.03.013 (2016).

39 Zhou, F. Molecular Mechanisms of IFN-γ to Up-Regulate MHC Class I Antigen Processing and Presentation. International Reviews of Immunology 28, 239–260, doi:10.1080/08830180902978120 (2009).

40 Dejardin, E. et al. Regulation of major histocompatibility complex class I expression by NF-κB-related proteins in breast cancer cells. Oncogene 16, 3299–3307, doi:10.1038/sj.onc.1201879 (1998).

41 Li, A. et al. Activating cGAS-STING pathway for the optimal effect of cancer immunotherapy. Journal of Hematology & Oncology 12, 35, doi:10.1186/s13045-019-0721-x (2019).

42 Thomson, D. W. et al. Discovery of GSK8612, a Highly Selective and Potent TBK1 Inhibitor. ACS Medicinal Chemistry Letters 10, 780–785, doi:10.1021/acsmedchemlett.9b00027 (2019).

43 Podolin, P. L. et al. Attenuation of murine collagen-induced arthritis by a novel, potent, selective small molecule inhibitor of IkappaB Kinase 2, TPCA-1 (2-[(aminocarbonyl)amino]-5-(4-fluorophenyl)-3-thiophenecarboxamide), occurs via reduction of proinflammatory cytokines and antigen-induced T cell Proliferation. J Pharmacol Exp Ther 312, 373–381, doi:10.1124/jpet.104.074484 (2005).

44 Burke, J. R. et al. BMS-345541 is a highly selective inhibitor of I kappa B kinase that binds at an allosteric site of the enzyme and blocks NF-kappa B-dependent transcription in mice. J Biol Chem 278, 1450–1456, doi:10.1074/jbc.M209677200 (2003).

45 Napoleon, J. V. et al. Small-molecule IKKβ activation modulator (IKAM) targets MAP3K1 and inhibits pancreatic tumor growth. Proceedings of the National Academy of Sciences 119, e2115071119. doi:doi:10.1073/pnas.2115071119 (2022).

46 Yeo, H., Lee, Y. H., Koh, D., Lim, Y. & Shin, S. Y. Chrysin Inhibits NF-κB-Dependent CCL5 Transcription by Targeting IκB Kinase in the Atopic Dermatitis-Like Inflammatory Microenvironment. Int J Mol Sci 21, doi:10.3390/ijms21197348 (2020).

47 Davis, M. I. et al. Comprehensive analysis of kinase inhibitor selectivity. Nat Biotechnol 29, 1046–1051, doi:10.1038/nbt.1990 (2011).

48 Price, D. H. P-TEFb, a cyclin-dependent kinase controlling elongation by RNA polymerase II. Mol Cell Biol 20, 2629–2634, doi:10.1128/mcb.20.8.2629-2634.2000 (2000).

49 Ni, Z. et al. P-TEFb is critical for the maturation of RNA polymerase II into productive elongation in vivo. Mol Cell Biol 28, 1161–1170, doi:10.1128/mcb.01859-07 (2008).

50 Chao, S.-H. et al. Flavopiridol inhibits P-TEFb and blocks HIV-1 replication. Journal of Biological Chemistry 275, 28345–28348 (2000).

51 McPartland, R. P., Wang, M. C., Bloch, A. & Weinfeld, H. Cytidine 5’-triphosphate synthetase as a target for inhibition by the antitumor agent 3-deazauridine. Cancer Res 34, 3107–3111 (1974).

52 Nilson, Kyle A. et al. THZ1 Reveals Roles for Cdk7 in Co-transcriptional Capping and Pausing. Molecular Cell 59, 576–587, doi:10.1016/j.molcel.2015.06.032 (2015).

53 King, H. M. et al. Aminopyrazole based CDK9 PROTAC sensitizes pancreatic cancer cells to venetoclax. Bioorg Med Chem Lett 43, 128061, doi:10.1016/j.bmcl.2021.128061 (2021).

54 Twyman-Saint Victor, C. et al. Radiation and dual checkpoint blockade activate non-redundant immune mechanisms in cancer. Nature 520, 373–377, doi:10.1038/nature14292 (2015).

55 Dexter, D. L. et al. Activity of a novel 4-quinolinecarboxylic acid, NSC 368390 [6-fluoro-2-(2’-fluoro-1,1’-biphenyl-4-yl)-3-methyl-4-quinolinecarb oxylic acid sodium salt], against experimental tumors. Cancer Res 45, 5563–5568 (1985).

56 Klotz, L. et al. Teriflunomide treatment for multiple sclerosis modulates T cell mitochondrial respiration with affinity-dependent effects. Science Translational Medicine 11, eaao5563. doi:doi:10.1126/scitranslmed.aao5563 (2019).

57 Fox, R. I. et al. Mechanism of action for leflunomide in rheumatoid arthritis. Clin Immunol 93, 198–208, doi:10.1006/clim.1999.4777 (1999).

58 Miller, A. E. An updated review of teriflunomide’s use in multiple sclerosis. Neurodegener Dis Manag 11, 387–409, doi:10.2217/nmt-2021-0014 (2021).

59 Shin, D. S. et al. Primary Resistance to PD-1 Blockade Mediated by JAK1/2 Mutations. Cancer Discov 7, 188–201, doi:10.1158/2159-8290.Cd-16-1223 (2017).

60 Dersh, D. et al. Genome-wide Screens Identify Lineage- and Tumor-Specific Genes Modulating MHC-I- and MHC-II-Restricted Immunosurveillance of Human Lymphomas. Immunity 54, 116–131.e110, doi:10.1016/j.immuni.2020.11.002 (2021).

61 Petroni, G., Buqué, A., Zitvogel, L., Kroemer, G. & Galluzzi, L. Immunomodulation by targeted anticancer agents. Cancer Cell 39, 310–345, doi:10.1016/j.ccell.2020.11.009(2021).

62 Keshet, R. et al. Targeting purine synthesis in ASS1-expressing tumors enhances the response to immune checkpoint inhibitors. Nature Cancer 1, 894–908, doi:10.1038/s43018-020-0106-7 (2020).

63 Olou, A. A., King, R. J., Yu, F. & Singh, P. K. MUC1 oncoprotein mitigates ER stress via CDA-mediated reprogramming of pyrimidine metabolism. Oncogene 39, 3381–3395, doi:10.1038/s41388-020-1225-4 (2020).

64 Dasgupta, A. et al. SIRT1-NOX4 signaling axis regulates cancer cachexia. J Exp Med 217, doi:10.1084/jem.20190745 (2020).

65 Shukla, S. K. et al. Silibinin-mediated metabolic reprogramming attenuates pancreatic cancerinduced cachexia and tumor growth. Oncotarget 6, 41146–41161, doi:10.18632/oncotarget.5843 (2015).

