## Supplementary figures and images for "DHODH inhibition enhances the efficacy of immune checkpoint blockade by increasing cancer cell antigen presentation"

### Supplementary Figure 1

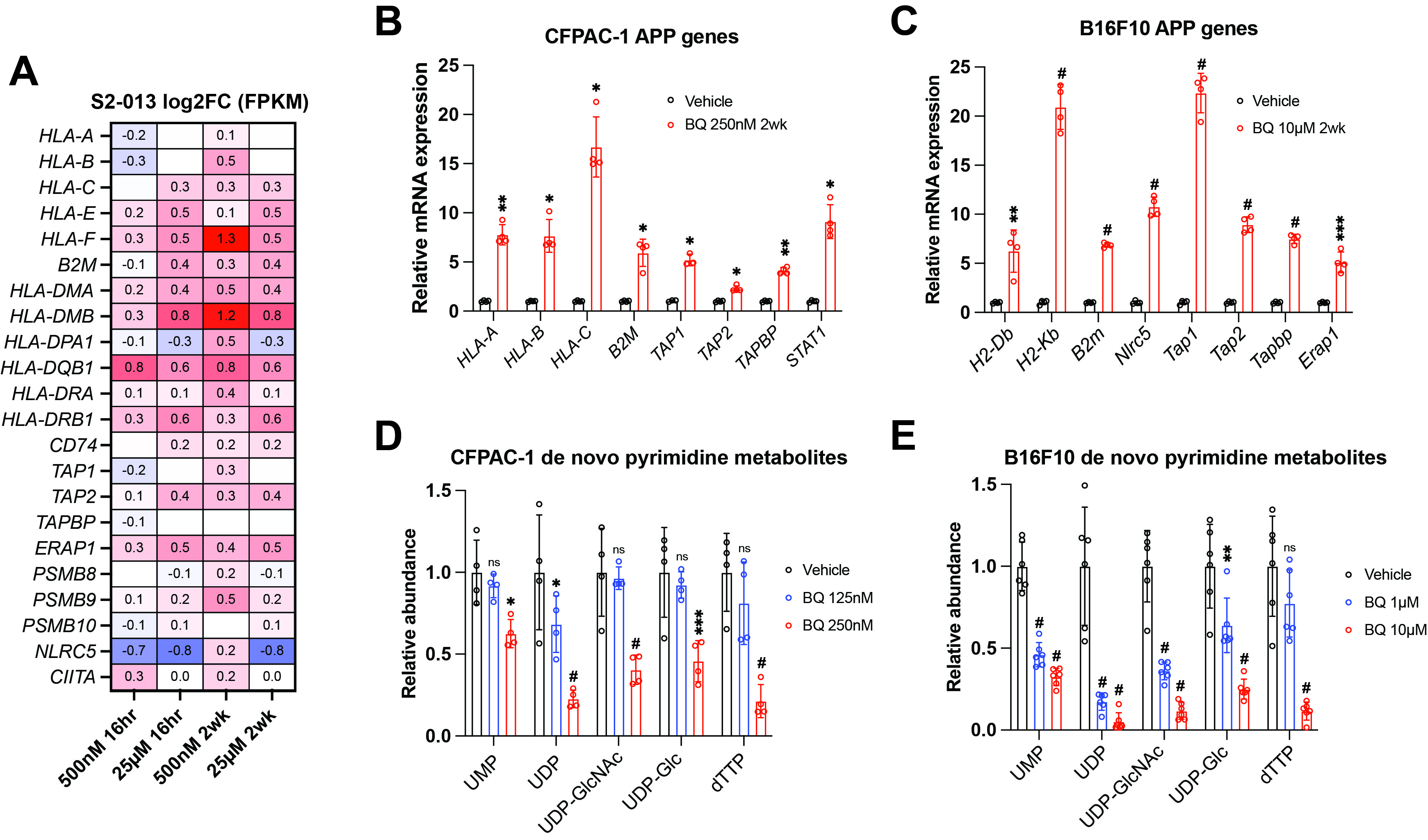

### Supplementary Figure 2

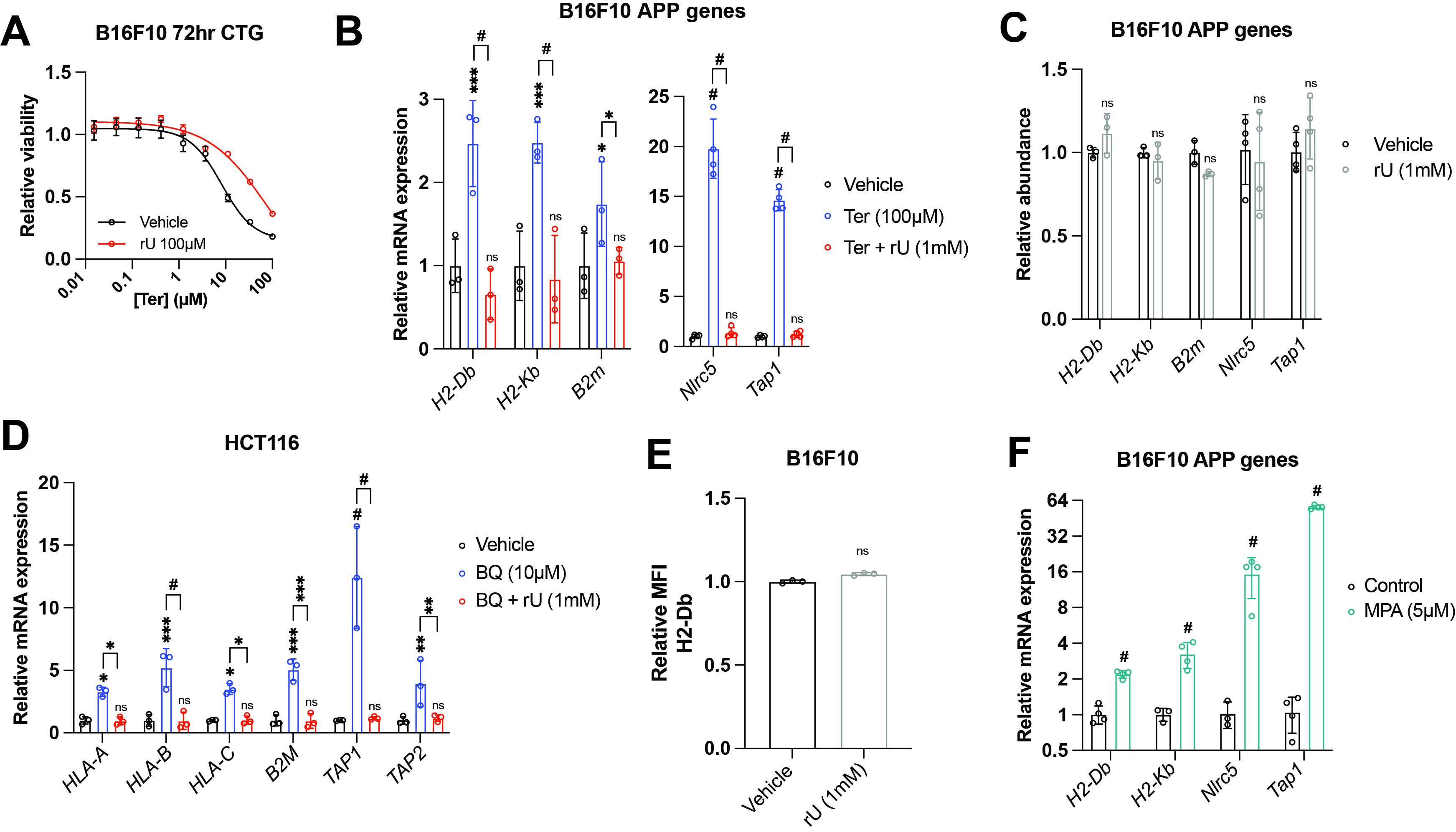

### Supplementary Figure 3

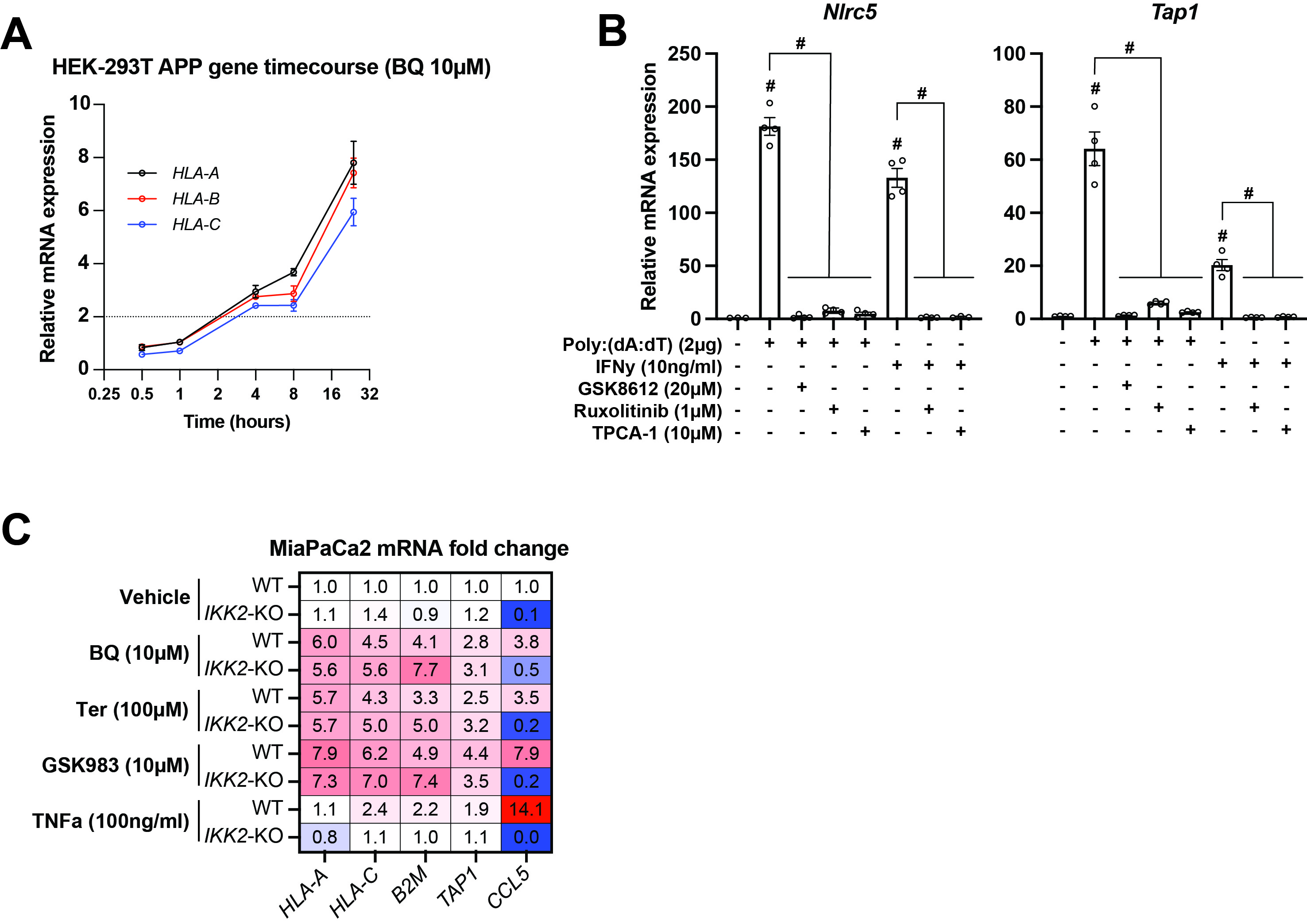

### Supplementary Figure 4

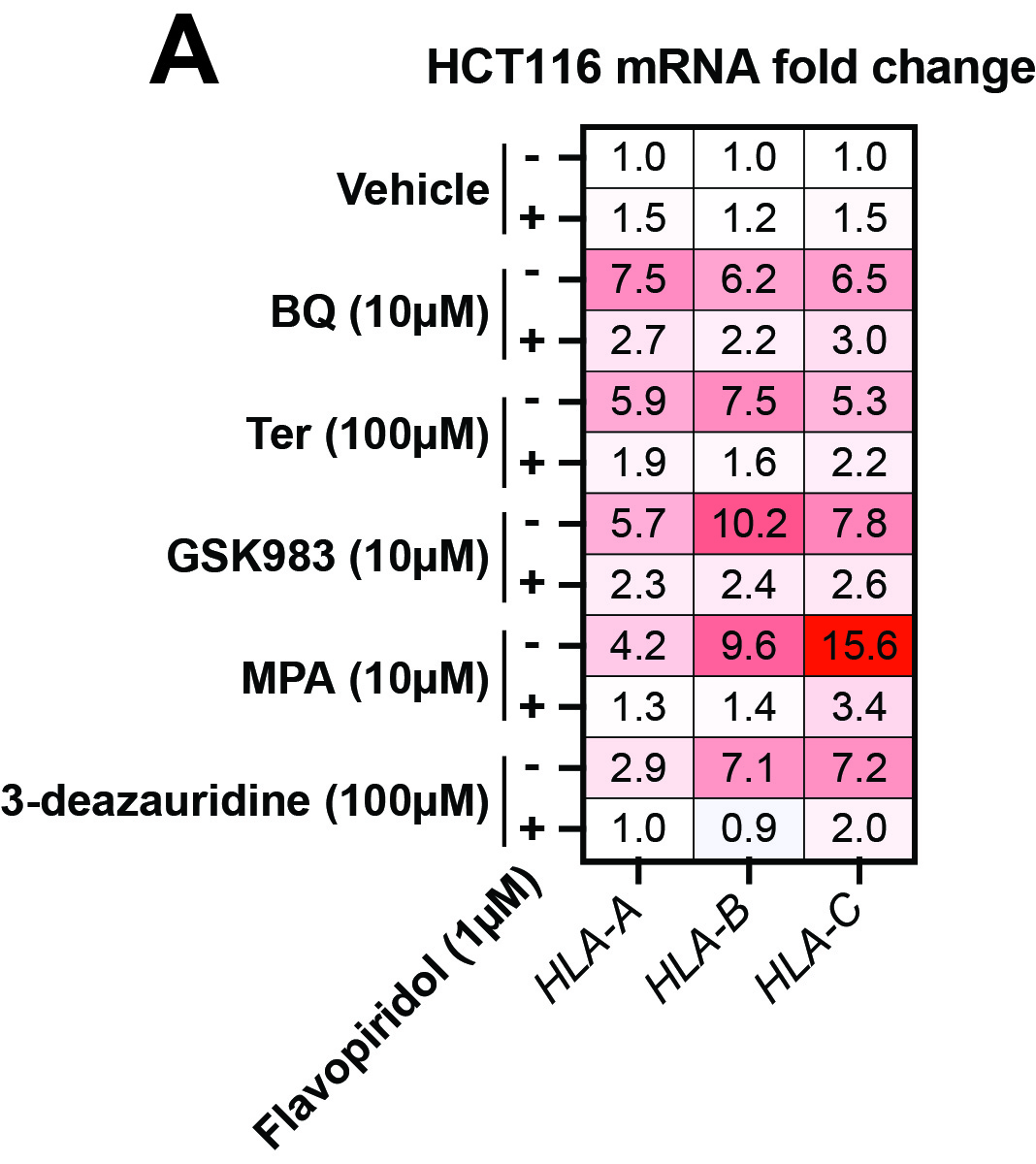

### Supplementary Figure 5

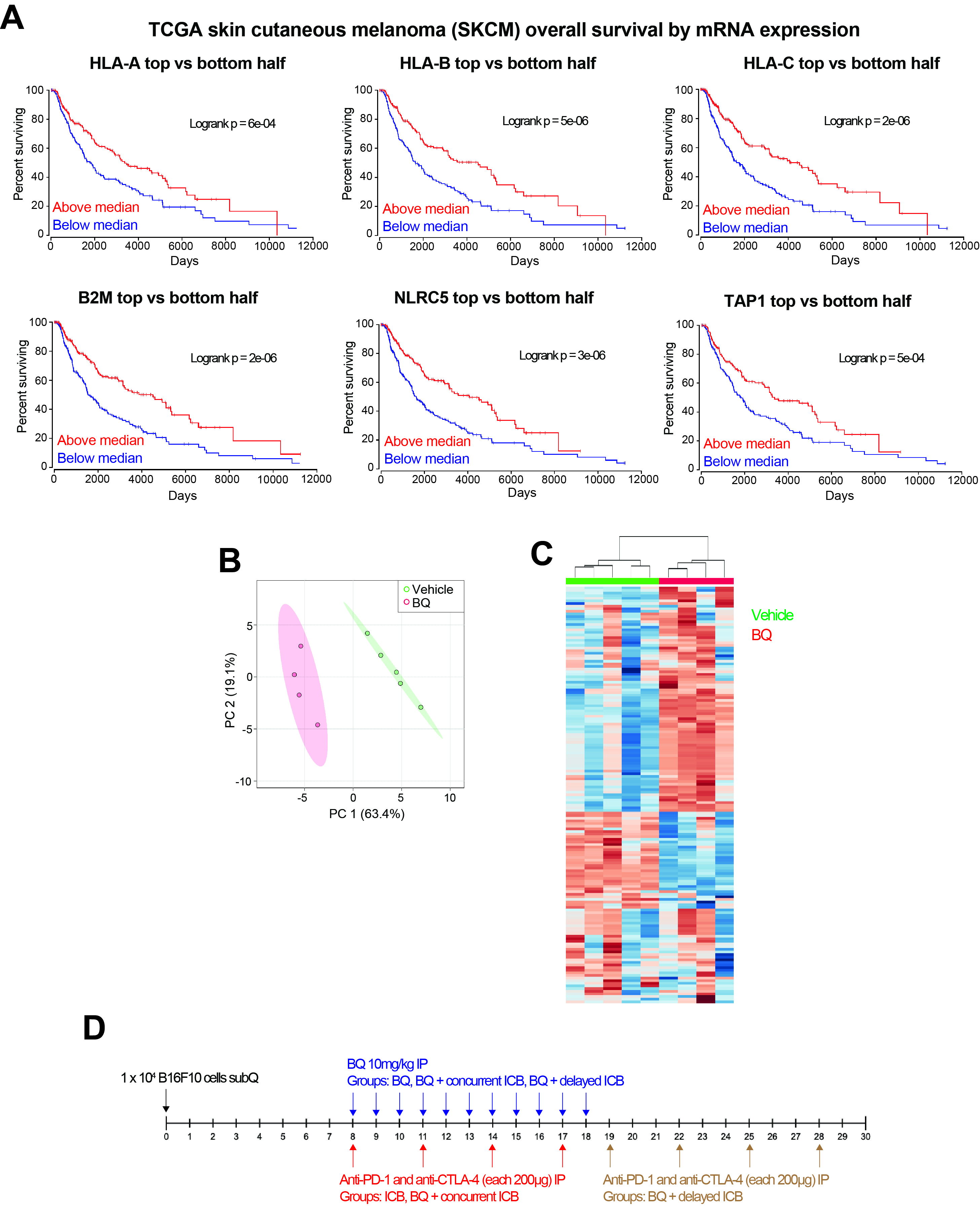
